# *mspms*: A Comprehensive R Package and Graphic Interface for Multiplex Substrate Profiling by Mass Spectrometry Analysis

**DOI:** 10.1101/2025.04.14.648679

**Authors:** Charlie Bayne, Brianna Hurysz, David J. Gonzalez, Anthony O’Donoghue

## Abstract

Multiplex Substrate Profiling by Mass Spectrometry (MSP-MS) is a powerful method for determining the substrate specificity of proteolytic enzymes, knowledge key for developing protease inhibitors, diagnostics, and protease-activated therapeutics. However, the complex datasets generated by MSP-MS pose significant analytical challenges. To address this, we developed *mspms*, a Bioconductor R package complemented by an intuitive graphical interface. *Mspms* streamlines MSP-MS data analysis by standardizing workflows for data preparation, processing, statistical analysis, and visualization. Designed for accessibility, it serves both advanced users via the R package and broader audiences through the web interface. We validated *mspms* by profiling the substrate specificity of four well-characterized cathepsins (A–D), demonstrating its ability to reliably capture expected substrate specificities. As the first publicly available platform for MSP-MS data analysis, *mspms* delivers comprehensive functionality, transparency, and ease of use, making it a valuable resource for the protease research community. Access to *mspms* is available through the Bioconductor project at https://bioconductor.org/packages/mspms, and a graphic interface is available at https://gonzalezlab.shinyapps.io/mspms_shiny/.

**Author Summary:** We developed mspms, an easy-to-use tool that helps researchers analyze data from a proteomics technique called Multiplex Substrate Profiling by Mass Spectrometry (MSP-MS). This software improves on previous methods of analyzing MSP-MS data, which required the user to navigate a confusing mix of R scripts, manual manipulation of spreadsheets, and third-party tools—an approach that was daunting for collaborators and new graduate students alike. Mspms streamlines the process, enabling faster, more reliable, and reproducible data analysis. We tested the tool using well-known proteases and found that it accurately identifies their known targets. As the first comprehensive tool for MSP-MS analysis, mspms makes this method approachable to a wider audience. It’s available for free through the Bioconductor project at https://bioconductor.org/packages/mspms, and a graphical interface is available at https://gonzalezlab.shinyapps.io/mspms_shiny/.

## Introduction

Proteases play crucial roles in a wide range of biological processes, from digestion and immunity to cancer and neurodegenerative diseases (1). Understanding the substrate specificity of these enzymes is essential for designing inhibitors, diagnostics, and protease-activated therapeutics (2). One of the most effective methods for determining protease substrate specificity is Multiplex Substrate Profiling by Mass Spectrometry (MSP-MS) (3). This technique involves incubating a rationally designed peptide library with a protease or protease-containing sample and using mass spectrometry to identify the resulting cleavage products, revealing the enzyme’s substrate preferences (4).

The data produced by MSP-MS are complex and multi-dimensional. Accurate interpretation of these results requires rigorous data analysis, encompassing multiple steps: preparing the data (identifying cleavage motifs, and positions), processing the data (data transformation, normalization, and imputation), statistical analysis, and data visualization. Historically, analysis of MSP-MS data has lacked dedicated analysis tools, leaving each researcher to analyze their data in an *ad-hoc* manner. This approach, while functional, results in an inherent lack of reproducibility. Inconsistent and irreproducible analysis pipelines have been noted to lead to grave problems in biological research (5), often a consequence of decentralized, error prone evolution of codebases as personnel transitions (6). Such absence of standardized, reproducible data analysis tools for MSP-MS has limited scientific progress by creating a barrier for collaboration across research groups.

To address these challenges, we developed *mspms*, an R package specifically designed for the robust, reproducible analysis of MSP-MS data. Through integration into the Bioconductor ecosystem (7), the *mspms* package adheres to best practices in software development and data analysis, offering a transparent and portable solution for processing complex datasets.

Recognizing that many users may not have programming experience, we complemented the R package with a user-friendly graphical interface, available both as a web application and for download. This interface allows researchers to perform key MSP-MS analysis steps—data preprocessing, normalization, statistical analysis, and visualization—without needing any R programming knowledge.

Here, we demonstrate the functionality of *mspms* by analyzing publicly available MSP-MS data for four well-characterized cathepsins, validating the package’s ability to accurately determine their substrate specificities. By offering comprehensive functionality, transparency, and user accessibility, *mspms* is positioned to be a valuable tool for the protease research community, streamlining the analysis of MSP-MS data while promoting reproducible research.

## Materials and Methods

### Data Used for Study

Raw data from a previously reported MSP-MS study was acquired from the MassIVE Repository (accession number MSV00008595) (8). In brief, these data were generated from a study that utilized a 228-member peptide library that was incubated with either 18.4 nM cathepsin A, 2.64 nM cathepsin B, 19.6 nM cathepsin C or 100 nM cathepsin D. The concentration of each peptide in the library was 0.5 μM. After incubation at 37°C for defined time points, the reaction was quenched by addition of 6.4 M guanidine hydrochloride. These samples were desalted with C18 spin columns, and ∼0.4 μg of each sample was subjected to LC-MS/MS analysis using an Ultimate 3000 HPLC and Q-Exactive mass spectrometer. LC and MS parameters were as previously reported (8).

### Upstream Proteomic Software

Peptides/proteins were identified and quantified using PEAKS Studio (9), Proteome Discoverer (10), and FragPipe (11). The database used in each search was the 228-member peptide library described previously (3) (Supplementary File 1).

### PEAKS Studio

Data from all .raw files were processed using PEAKS Studio v8.5 software, using a customized template (Supplementary File 2). For each sample experiment specific parameters were set as follows: Q-Exactive instrument, HCD fragmentation, no enzyme. Scans were merged with a retention time window of 0.8 min, and precursor m/z error tolerance of 10 ppm. Precursor mass was corrected. Scans were filtered to include retention time between 0 and 95 min with a precursor mass tolerance of 10 ppm. For identification, a precursor mass tolerance of 20 ppm using monoisotopic mass and a fragmentation ion of 0.01 Da was specified. No PTMs were included in the search. FDR was estimated using decoy-fusion strategy. Label free quantification was performed with a mass error tolerance of 9 ppm, and retention time shift tolerance of 3 minutes. Replicate samples were added to new groups.

The peaks_protein-peptides-lfq.csv file was prepared by navigating to the quantification options setting the normalization factor to “No normalization”, changing peptide filters to include all peptides (quality ≥ 0, Avg.Area ≥ 0, Peptide ID Count ≥0, charge +1 - +10, and at least 1 confident sample). Protein filters were changed so no filtering occurs (Significance ≥0). Data was then exported as the peptides-lfq.csv file.

Data in figures is derived from the PEAKS software, unless otherwise specified.

### Proteome Discoverer

Data from all .raw files was also processed using Proteome Discoverer V 2.5.0.400 using a customized processing and consensus workflow. (Supplemental Files 3 and 4). Briefly, min precursor mass was specified as 350 Da, max precursor mass was specified as 5000. Enzyme was set to be unspecific, with a min peptide length of 5, and max peptide length of 14. Precursor mass tolerance was set to 10 ppm, and fragment mass tolerance was set to 0.6. Percolator Target FDR was set to 0.01.

### FragPipe

FragPipe V22.0. MSFragger version 4.1, IonQuant version 1.10.27, and Python version 3.9.13 were used to process all .raw files with a customized analysis workflow derived from the MBR- LFQ workflow template (Supplementary File 5). Briefly, decoys were added to the peptide library database, cleavages were set to nonspecific; peptide length was set at 5-14, and 350 Da to 5000 Da, match between runs was enabled, and top runs was set to 3 (as there were 4 biological replicates in each group except for cathepsin D at time zero).

### Preprocessing

To prepare MSP-MS data for analysis, *mspms* preprocesses an exported file from the user’s proteome software of choice. In this process, the data is converted to a standardized format and loaded as a QFeatures object (12) containing a SummarizedExperiment (13) object named “peptides”, which contains the detected peptide intensities. Cleavage motifs of a user specified length to the left and right of the scissile bond are calculated, and the numerical location of the cleavage site (via reference to the member of the library it was derived from) is determined.

These peptide centric features are then loaded as the rowData corresponding to the QFeatures object. The colData composing the QFeatures experiment contains sample metadata describing the experiment and must include descriptors core to every MSP-MS experiment: “group”, “condition”, and “time”.

### Data Processing

Peptide values are subjected to log_2_ transformation followed by a median centered normalization utilizing the center.median method. Due to the left-censored nature of MSP-MS data, imputation is subsequently performed using the QRILC method. Lastly, the data is reverse log_2_ transformed. All data manipulation is performed using MScoreutils (14). Data resulting from each step of data processing is stored within the resulting QFeatures object as SummarizedExperiment objects named “log2_peptides”, “log2_peptides_norm”, “log2_peptides_norm_imputed”, and “peptides_norm” respectively.

### Statistics

*Mspms* provides two methods for performing statistics. The first method calculates the log_2_ fold change relative to the user specified denominator in colData and then performs pairwise t-tests with FDR p adjustment as implemented in the Rstatix (15) package using the peptides_norm values. The second method utilizes the limma R package (16), which is widely used for analyzing high-dimensional data, particularly in genomics and proteomics. Limma applies linear models to assess differential expression across different conditions and handles complex experimental designs, accounting for factors such as batch effects, repeated measures, and covariates. It also offers robustness by using empirical Bayes methods to stabilize variance estimates, making it particularly useful when sample sizes are small or when there is high variability in the data. In the present study, t-test results are reported for maximum interpretability, though similar outcomes were obtained with both methods. We note that the limma package is significantly faster and provides greater flexibility for handling more complex experimental designs.

By default, significant peptides are denotated as having a p.adj < 0.05 and log_2_ fold change > 3 relative to time 0.

### Visualizations

*Mspms* supports ggplot2 (17) based plotting of several types of static visualizations including quality control, PCA, volcano, time course, and iceLogo plots. Interactive heatmaps are plotted using the plotly (18) based heatmaply (19) library .

### iceLogo Analysis

iceLogo (20) analysis calculates the chance of occurrence (p-value) of every amino acid surrounding the scissile bond in the experimental data relative to a user-defined reference set of possible scissile bond locations. We implemented this approach in R, using the previously described Java implementation as a reference. The underlying logic is described below. First, a count of the number of times an amino acid at each position is calculated for an experimental and reference set.

Then the frequency of each amino acid at each position is calculated.

Then the standard deviation (𝜎) is calculated using the frequency of an amino acid in the reference set (𝑓%)

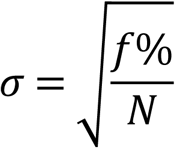

These calculated standard deviations are subsequently used to calculate significances by conversion to p-values using the Wichura algorithm. Only p values ≤ to the user specified p value threshold are retained for subsequent visualization.

The height of each amino acid at each position is then visualized using the ggseqlogo R package (21) using the user’s choice of percent change or fold change to represent the height of each amino acid letter.

The percent change (𝑃𝐶) is calculated from the experimental set frequency (𝐹^+^) and reference set frequency (𝐹^―^) as follows:

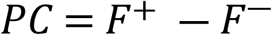

The fold change (𝐹𝐶) is calculated from the experimental set frequency (𝐹^+^) and reference set frequency (𝐹^―^) as follows:

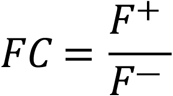

In the event that the fold change is smaller than one, it is transformed into the converted fold change (𝐹𝐶𝑐𝑜𝑛) in order to allow the comparison of height with positively regulated amino acids.

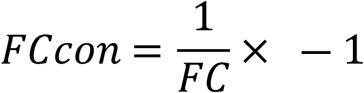

If only one amino acid is found to be significant at a given position and the calculated amino acid size is infinite, the height of the amino acid is represented as the maximal height that can be visualized in the iceLogo plot.

If several amino acids are found to be significant at a given position and all have infinite calculated amino acid sizes, the height of the amino acids combined is represented as the maximal height that can be visualized in the iceLogo plot.

### Report Generation

*Mspms* supports the production of a generic *mspms* report (Supplementary File 6). This function produces a generic self-contained .html report with embedded downloadable data frames (containing normalized data and results of statistics), and figures. This report is produced by leveraging the *mspms* R package inside of a parameterized rmarkdown (22) template incorporating the downloadthis (23) package.

### Helper Functions

Only a subset of functions are exported to the user in order to maintain an intuitive application programming interface. Helper functions can be found in the helper_functions.R file named corresponding to the type of functions it assists with.

### Graphic Interface

A graphic interface to the *mspms* R package is implemented using the R shiny framework (24). This interface is accessible on the web at https://gonzalezlab.shinyapps.io/mspms_shiny/ or downloadable from https://github.com/baynec2/mspms-shiny/.

## Results

The *mspms* R package was developed to provide a dedicated tool to analyze MSP-MS data, focusing on reproducibility, ease of use, and robustness. It includes modular functions to handle key steps generalizable to any MSP-MS analysis: data preparation, processing, statistical analysis, and visualization (Figure 1).

**Figure 1.**
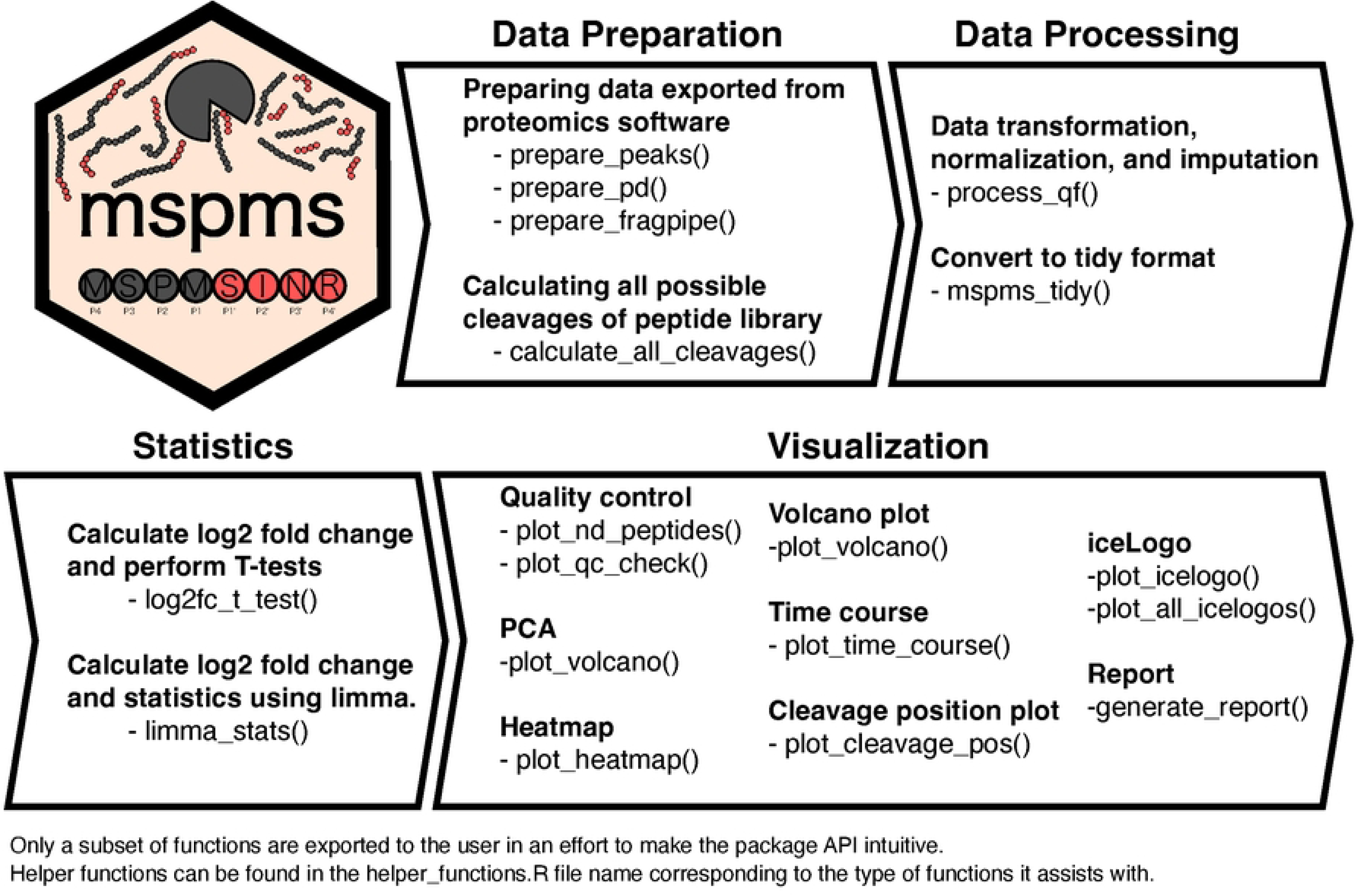
Overview of the *mspms* R package and MSP-MS profiling of cathepsin proteases. Schematic of the functions contained within the *mspms* R package.

### Data Quality Evaluation

To assess the quality of the MSP-MS data from the cathepsin A, B, C, and D experiments, we applied the quality control functions of *mspms*. We found that over 90% of the full-length peptide library was detected in all samples at time zero (T0), and more than 95% of the library, including cleavage products, was detected across the dataset (Supplementary Figure 4A). Only five peptides from the library were consistently missing across all samples, suggesting high-quality data with minimal loss (Supplementary Figure 4B).

### Evaluation of Global Data Patterns

Next, we examined global patterns in the dataset using principal component analysis (PCA) and unsupervised hierarchical clustering. PCA demonstrated tight clustering of replicates within each experimental group (condition and timepoint), as shown by the 95% confidence intervals surrounding each group (Figure 2A). Near-perfect clustering of replicates from identical conditions was observed, indicating high experimental consistency. Differential peptide abundance between groups was evident, supporting distinct activity for each cathepsin over time (Figure 2B).

**Figure 2.**
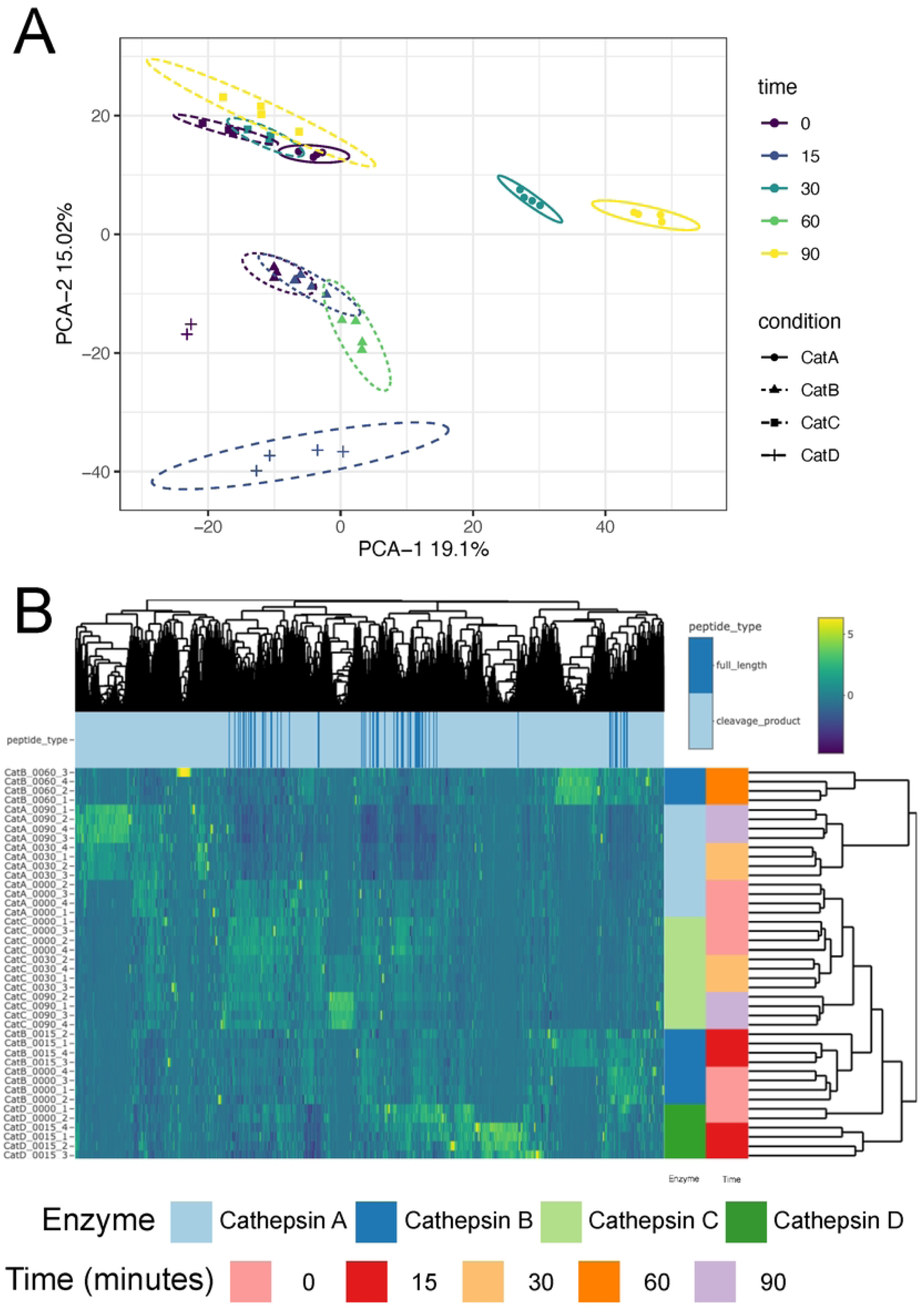
Global visualization of MSP-MS data. (A) Principal component analysis displaying PC1 and PC2. Samples are colored by time, while the shape and line type show the type of cathepsin with eclipses representing the 95% confidence interval. (C) Heatmap showing the results of the experiment as clustered using unsupervised hierarchical clustering. Rows of the heatmap represent the samples while columns represent the peptides. Color of the heatmap cells represent the normalized, column centered, and scaled values. Colored bars to the right of the heatmap indicate the cathepsin and time of the samples in each row. Colored bars corresponding to each peptide in the columns display whether the corresponding peptide is a full-length peptide belonging to the 228-member peptide library (non-cleaved, dark blue) or a cleavage product (cleaved, blue)

### Significant Peptide Changes and Cleavage Position Preferences

We analyzed significant peptide differences for each cathepsin relative to T0 using t-tests and log_2_ fold change calculations. The results, visualized as volcano plots, revealed numerous significantly upregulated peptides (log_2_ fold change ≥ 3, p.adj ≤ 0.05) at various timepoints following incubation (Figure 3B). The number of significantly different peptides increased progressively with time for each cathepsin, highlighting the dynamic substrate cleavage behavior (Supplementary Figure 5).

**Figure 3.**
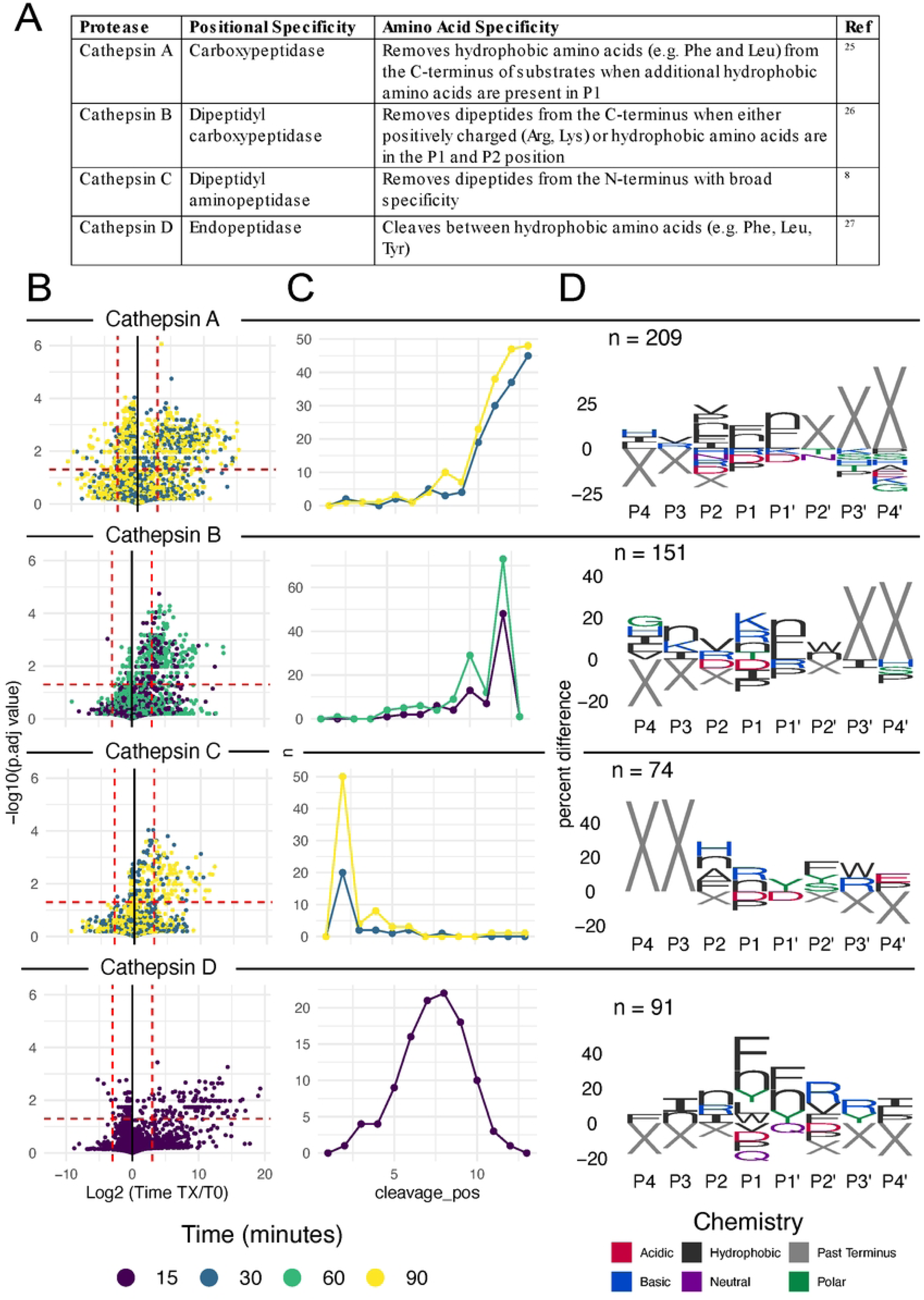
Differentially abundant peptide cleavages over time. (A) Summarized substrate specificities for cathepsin A-D as reported in the literature. (B) Volcano plots displaying the log_2_-fold change of the timepoint as indicated by color relative to and -log10 FDR corrected p values for each cathepsin. (C) Plot showing the number of significant cleavage events at each position of the peptide library (as defined as having a log_2_ fold change ≥ 3 and FDR adjusted p values ≤ 0.05) (D) IceLogo plots as implemented in the *mspms* package. Amino acid residues (with X representing positions past the terminus) four positions to the left and right of the cleavage site are displayed. Only residues with significantly higher proportions relative to the proportion of all possible cleavage sequences present in the initial peptide library (pval ≤ 0.05) are shown, with the height representing the percentage differences.

To evaluate protease activity relative to reported substate specificities (Figure 3A), we investigated cleavage site preferences within the 14-mer peptides. Cathepsin A showed clear carboxypeptidase activity through the high number of cleavage sites at the C-terminus (Figure 3C) and overrepresentation of X (corresponding to no amino acid) at P2’, P3’, and P4’ (Figure 3D). Cathepsin B displayed dipeptidyl carboxypeptidase activity, with most cleavage sites occurring at position 12 and a lesser amount at position 10, suggesting sequential removal of dipeptides from the C-terminus (Figure 3C). The enrichment of X at P3’ and P4’ evident in the iceLogo plot further supported this dipeptidyl carboxypeptidase activity (Figure 3D).

Assessment of cathepsin C revealed a high number of cleavage sites at position 2, followed by a smaller number at position 4, suggesting the sequential removal of dipeptides from the N-terminus (Figure 3C). An overrepresentation of X at P4 and P3 in the iceLogo also confirmed the dipeptidyl aminopeptidase activity (Figure 3D). Lastly, cathepsin D showed endopeptidase activity as observable by the Gaussian peak centered around position 8 of the cleavage position plots (Figure 3C). The iceLogo plot showed that X was not enriched at any of the sites from P4 to P4′ further validating this endopeptidase activity (Figure 3D).

### Amino Acid Preferences

To visualize the amino acid preferences at cleavage sites, we performed an iceLogo analysis using *mspms*, focusing on the eight positions surrounding the cleavage site (P4 to P4′).

Cathepsin A showed a preference for the removal of hydrophobic amino acids (such as Phe and Leu) from the C-terminus of substrates, when additional hydrophobic residues occupy the P1 position (Figure 3D). Cathepsin B favored substrates with positively charged (Arg, Lys) or hydrophobic residues in P1 and P2 (Figure 3D). Cathepsin C showed a limited preference at the P1 and P1′ position (Figure 3D). Cathepsin D showed a preference for Phe, Tyr, and norleucine at the P1 and P1′ position (Figure 3D).

### Comparison of Results Across Different Upstream Proteomics Software

To demonstrate the compatibility of MSP-MS with a range of upstream proteomics software, we analyzed MSP-MS data using PEAKS Studio, Proteome Discoverer (PD), and FragPipe separately, comparing their outputs (Supplementary Figure 1).

Processed data corresponding to peptides detected across all three approaches correlated well, with R^2^ values ranging from 0.70 (FragPipe to PD) to 0.81 (FragPipe to PEAKS) (Supplementary Figure 1A). When assessing peptides identified as significantly different from time zero, approximately 30% of significant peptides from cathepsins A, B and C were shared across all three platforms (Supplementary Figure 1B). For cathepsin D, the only endopeptidase in our study, there was only 17% agreement across the three software tools. However, an increase in shared peptide identities was observed between FragPipe and PEAKS for cathepsin D (27%) compared to cathepsins A–C (∼10%) (Supplementary Figure 1B).

Positional specificity, as evaluated through cleavage position plots, was highly consistent for cathepsins A, B and C across all tools. However, PD showed limited ability to capture the expected endopeptidase activity of cathepsin D, while FragPipe results displayed more N-terminal cleavage than observed in PEAKS data (Supplementary Figure 1C).

The specificity profiles at positions P4–P4′, evaluated using iceLogos, were comparable across all tools, with subtle differences depending on the software. Major discrepancies were observed in the cathepsin D results, consistent with findings from cleavage position plots (Supplementary Figure 2).

Since cathepsin D was the only endopeptidase included in this study, we hypothesized that the observed discrepancies arose from differences in each tool’s ability to detect shorter peptides, which are more commonly generated by endopeptidases. Analyzing the distribution of significant peptides by length revealed that PD systematically detected fewer peptides shorter than eight amino acids compared to PEAKS and FragPipe (Supplementary Figure 3). This likely explains PD’s poorer performance on cathepsin D compared to the other tools.

## Discussion

Before the development of the *mspms* package, MSP-MS data analysis relied heavily on ad hoc developed R scripts, which posed significant challenges. These scripts were fragmented, poorly documented, and difficult to adapt, making reproducibility a concern. Researchers without specialized programming skills struggled with customizing these workflows to accommodate different experimental designs, limiting the broader utility of MSP-MS data analysis.

Furthermore, only data exported from the proteomic search engine PEAKS was compatible, hindering usability across different research groups.

The *mspms* R package effectively addresses key limitations in MSP-MS data analysis through its modular, reproducible, user-friendly approach, and compatibility with a wide range of proteomics software. It provides self-contained functions for data preparation, processing, statistical analysis, and visualization, ensuring ease of maintenance, extensibility, and usability.

One significant feature is the integration of functionality from the widely cited iceLogo tool within R, which allows for the analysis of nonstandard amino acids, such as norleucine, and positions marked by “X.” Additionally, *mspms* includes a graphical user interface, accessible both online and via local download, enabling researchers without R programming experience to leverage its core functionalities. Moreover, *mspms* integrates smoothly with the Bioconductor ecosystem. By employing established S4 classes internally, it offers R users the flexibility to tap into Bioconductor’s extensive analytical resources, further enabling advanced data exploration, statistical analysis, and visualizations. This makes *mspms* a versatile and adaptable tool that meets the diverse needs of the protease research community.

In the present study, we applied *mspms* to profile the substrate specificity of four well-characterized cathepsin proteases, namely cathepsin A, B, C, and D. In doing so, we demonstrate features of *mspms* that are broadly applicable to any MSP-MS experiment, while rigorously benchmarking the software by evaluating its ability to detect substrate specificities accepted to be the ground truth.

A critical but often overlooked step in proteomic analysis is conducting a thorough quality control assessment to ensure data quality is sufficient for drawing biologically meaningful conclusions (25). Since each MSP-MS experiment is based on a known peptide library, an effective quality control measure involves evaluating the percentage of the un-cleaved peptide library detected in each sample. Ideally, 100% of the un-cleaved peptide library should be detectable at T0; however, due to limitations in mass spectrometry performance, this is rarely achieved in practice. When we applied this quality control check to our cathepsin experiment, we observed no indication of data quality issues, confirming the reliability of our results.

The next step of an MSP-MS analysis is to explore global data patterns present in the data. This approach allows users the ability to evaluate the data as a whole, determine whether the experiment was technically successful (by verifying that the positive and negative controls performed as expected), and identify interesting patterns. *Mspms* allows the user to easily create PCA and interactive heatmap plots. In our cathepsin MSP-MS experiment, PCA and heatmap analyses revealed tight clustering among replicate conditions, indicating minimal variability across biological replicates. Moreover, distinct clustering of different experimental groups based on cathepsin type and time point highlighted the unique substrate specificities of each enzyme. These visualizations underscore the reliability of the experiment and the capacity of *mspms* to facilitate robust, comprehensive data exploration.

Once the MSP-MS experiment is confirmed as a technical success, the main objective— determining the enzymes’ substrate specificity—is readily achievable using the *mspms* tool. First, *mspms* computes the log₂ fold change and FDR-corrected p-values from t-tests conducted on normalized and imputed intensity values. The specific features of significantly altered peptides are then examined via cleavage location plots, which illustrate the enzyme’s positional specificity, and iceLogo plots, which reveal amino acid preferences.

When applied to the cathepsin data, our analysis near perfectly captured previously reported substrate specificities for each of the enzymes profiled. We show that:

1. Cathepsin A is a carboxypeptidase that preferentially removes hydrophobic amino acids (such as Phe and Leu) from the C-terminus of substrates, especially when additional hydrophobic residues occupy the P1 position, as previously reported (26).
2. Cathepsin B is a dipeptidyl carboxypeptidase that cleaves dipeptides from the C-terminus, favoring substrates with positively charged (Arg, Lys) or hydrophobic residues in P1 and P2, as previously reported (27).
3. Cathepsin C functions as a dipeptidyl aminopeptidase, cleaving dipeptides from the N-terminus with broad specificity, as previously reported (8).
4. Cathepsin D is an endopeptidase that cleaves between hydrophobic amino acids, including Phe, Leu, and Tyr, as previously reported (28).

By validating the specific activities of these cathepsins, *mspms* confirms its capability to accurately identify expected substrate specificities, establishing its value as a powerful tool in protease research. Beyond the enzymes evaluated in this study, the package’s modular and flexible design enables it to be readily applied to analyze the substrate specificity of virtually any protease mixture, allowing for diverse applications across the protease field.

Moreover, *mspms* is built to integrate seamlessly with future advancements in MSP-MS assays. As peptide synthesis becomes more cost-effective and mass spectrometer technology advances, it will be feasible to expand the MSP-MS assay by incorporating significantly larger peptide libraries than the current 228-member set. The ability to support any peptide library, coupled with its reproducible workflows and user-friendly features, secures *mspms* as an enduring and valuable asset to the protease research community.

To enhance accessibility, MSP-MS is designed to be compatible with three major proteome search engines: PEAKS Studio, Proteome Discoverer (PD), and FragPipe. Compatibility was validated by independently analyzing MSP-MS data using each software tool. Substrate specificity interpretations for cathepsins A, B, and C were remarkably consistent across all platforms.

For cathepsin D, both PEAKS and FragPipe effectively detected endopeptidase activity, but FragPipe identified a higher frequency of significant N-terminal cleavages relative to PEAKS. Determining which profile is more biologically accurate would require further validation using orthogonal assays. Proteome Discoverer, in contrast, struggled to convincingly identify cathepsin D’s endopeptidase activity, likely due to its reduced sensitivity for shorter peptides relative to PEAKS and FragPipe.

If endopeptidase activity is a primary focus, we recommend using PEAKS Studio or Fragpipe, which provided the best detection of cathepsin D activity in our tests. We note that FragPipe is an attractive option, particularly as it is an open-source tool freely available for academic use and demonstrated analysis speeds at least an order of magnitude faster than the paid software solutions. We caution against using Proteome Discoverer with the search settings applied in this study unless the parameters are further optimized, which may be able to increase the ability to detect smaller peptides. If endopeptidase activity is not of interest, our results indicate that all three software platforms perform comparably.

## Conclusion

In summary, *mspms* streamlines MSP-MS data analysis, providing a reliable, reproducible, and adaptable platform for protease substrate profiling. The combination of its powerful analytical capabilities and intuitive design enables researchers to extract biologically meaningful insights from complex datasets with minimal technical barriers. Given its flexibility and broad applicability, *mspms* is positioned to become a standard tool in protease research, offering significant advancements in the study of proteolytic enzymes and their roles in health and disease.

## Availability of Source Code and Requirements

- Project name: mspms
- Operating system(s): Linux, macOS, Windows
- Programming language: R
- Other requirements: R 4.4.0, QFeatures, SummarizedExperiment, magrittr, rlang, dplyr, purrr, stats, tidyr, stringr, ggplot2, ggseqlogo, heatmaply, readr, rstatix, tibble, ggpubr.
- License: MIT
- Bioconductor home page: https://bioconductor.org/packages/mspms
- GitHub home page: https://github.com/baynec2/mspms
- Shiny app instance: https://gonzalezlab.shinyapps.io/mspms_shiny/
- Shiny app repository: https://github.com/baynec2/mspms-shiny
- Vignette : https://bioconductor.org/packages/devel/bioc/vignettes/mspms/inst/doc/mspms_vignette.html
- Manuscript repository: https://github.com/baynec2/mspms-manuscript.

## Availability of Supporting Data and Materials

All data used to build this manuscript can be found in the GitHub repository for the manuscript (https://github.com/baynec2/mspms_manuscript). Mass spectrometry data in. raw format is available from MassIVE Repository under accession number MSV00008595.

## Supplementary Figure Legends

**Supplementary Figure 1. Comparison of Results from Upstream Proteomic Software Compatible with *mspms*.**

(A) Correlation analysis of shared peptides detected by PEAKS Studio, Proteome Discoverer (PD), and FragPipe. (B) Venn diagram showing peptides identified as significantly different by each software tool. (C) Cleavage site plots illustrating peptide cleavage patterns across comparisons made using the three tools.

**Supplementary Figure 2. Comparison of IceLogos from Upstream Proteomic Software Compatible with *mspms*.**

IceLogo generated using peptide data exported from PEAKS Studio, Proteome Discoverer (PD), and FragPipe.

**Supplementary Figure 3. Comparison of Detected Peptide Lengths Using Different Upstream Proteomic Software Compatible with *mspms*.**

(A) Count of significant peptides detected by length, categorized by the upstream proteomic software tool used.

(B) Count of total detected peptides by length categorized by the upstream proteomic software tool used.

**Supplementary Figure 4. Quality Control Evaluation.**

(A) Histogram displaying the count of samples for each sample grouping at time 0 as a function of the percentage of undetected full-length and cleavage product peptides mapping to the 228-peptide library. (B) Percent of samples that indicated member of the 228-peptide library was undetected in when considering only full-length or cleavage product peptides.

**Supplementary Figure 5. Number of Significant Differences Relative to T0 as a Function of MSP-MS Incubation Time.** The number of significantly enriched peptides relative to time 0 as shown per duration of time incubated with the indicated cathepsin.

**Supplementary Figure 6. IceLogo Analyses with Extended Cleavage Motifs.** IceLogo analysis of peptide cleavage motifs containing the 6 amino acids before and after the significantly enriched peptides with detected cleavage sites relative to all possible within the background of the 228-peptide library used for the experiment.

**Supplementary Figure 7. Screenshots of *mspms* graphic interface.**

(A) About page. (B) File upload page. Once files are uploaded, subsequent pages become available to the user. (C) Processed data page containing normalized and imputed data. (D) Page containing quality control plots. (E) Stats page containing results of log_2_ fold change and FDR corrected t-tests relative to time 0 for each condition. User selected peptides can be interactively plotted with either the normalized imputed data, or the raw intensities. (F) DataViz page containing PCA, interactive heatmap, volcano plot, or iceLogo plots. (G) Page containing button to generate a self-contained *mspms* html report.

## Supplementary Files

*Supplemental files can be found in the GitHub repository for this manuscript.*

**Supplementary File 1. Peptide library fasta file used as proteomics search database.**

**Supplementary File 2. Peptide library .csv file corresponding to Supplementary file 1.**

**Supplementary File 3. Screenshot of parameters used in PEAKS studio search.**

**Supplementary File 4. Proteome Discoverer analysis template used for *mspms* analysis.**

**Supplementary File 5. Proteome Discoverer consensus template used for *mspms* analysis.**

**Supplementary File 6. Fragpipe workflow used for *mspms* analysis.**

**Supplementary File 7. Generic *mspms* .html report for cathepsin A-D data.**

## Abbreviations

MSP-MS: Multiplex Substrate Profiling by Mass Spectrometry.
T0: Time zero
PCA: Principal Component Analysis.
PC1: Principal component 1.
PC2: Principal component 2.
FDR: False Discovery Rate.
PD: Proteome Discoverer.

## Author contributions

C.B wrote the R package, shiny app, documentation, and manuscript; B.H made substantial contributions to the conception and design of the work, while D.J.G. and A.J.O provided funding, oversaw the project and provided contributions to the conception and design of the work. All authors edited and approved the final version of the manuscript.

## Acknowledgements

We would like to acknowledge Dr. Zhenze Jiang for his initial conception of previous R scripts, as well as Dr. Lawrence Liu and Dr. Michael Yoon for maintaining and updating the scripts. We also thank Dr. Jiang and Dr. Yoon for running the MSP-MS experiment analyzed in this manuscript. Finally, we would like to acknowledge Diego F. Trujillo for suggestions to improve *mspms* and for educating new users how to perform the data analysis.

## References

1. López-Otín C, Bond JS. Proteases: Multifunctional Enzymes in Life and Disease. Journal of Biological Chemistry. 2008 Nov;283(45):30433–7.

2. Leung D, Abbenante G, Fairlie DP. Protease Inhibitors: Current Status and Future Prospects. J Med Chem. 2000 Feb 1;43(3):305–41.

3. Rohweder PJ, Jiang Z, Hurysz BM, O’Donoghue AJ, Craik CS. Multiplex substrate profiling by mass spectrometry for proteases. In: Methods in Enzymology [Internet]. Elsevier; 2023 [cited 2024 Apr 17]. p. 375–411. Available from: https://linkinghub.elsevier.com/retrieve/pii/S0076687922003901

4. O’Donoghue AJ, Eroy-Reveles AA, Knudsen GM, Ingram J, Zhou M, Statnekov JB, et al. Global identification of peptidase specificity by multiplex substrate profiling. Nat Methods. 2012 Nov;9(11):1095–100.

5. Miller G. A Scientist’s Nightmare: Software Problem Leads to Five Retractions. Science. 2006 Dec 22;314(5807):1856–7.

6. Casadevall A, Steen RG, Fang FC. Sources of error in the retracted scientific literature. FASEB j. 2014 Sep;28(9):3847–55.

7. Gentleman RC, Carey VJ, Bates DM, Bolstad B, Dettling M, Dudoit S, et al. Bioconductor: open software development for computational biology and bioinformatics. Genome Biology. 2004;

8. Jiang Z, Lietz CB, Podvin S, Yoon MC, Toneff T, Hook V, et al. Differential Neuropeptidomes of Dense Core Secretory Vesicles (DCSV) Produced at Intravesicular and Extracellular pH Conditions by Proteolytic Processing. ACS Chem Neurosci. 2021 Jul 7;12(13):2385–98.

9. Ma B, Zhang K, Hendrie C, Liang C, Li M, Doherty-Kirby A, et al. PEAKS: powerful software for peptide *de novo* sequencing by tandem mass spectrometry. Rapid Comm Mass Spectrometry. 2003 Oct 30;17(20):2337–42.

10. Orsburn BC. Proteome Discoverer—A Community Enhanced Data Processing Suite for Protein Informatics. Proteomes. 2021 Mar 23;9(1):15.

11. Yu F, Haynes SE, Nesvizhskii AI. IonQuant Enables Accurate and Sensitive Label-Free Quantification With FDR-Controlled Match-Between-Runs. Molecular & Cellular Proteomics. 2021;20:100077.

12. Gatto L, Vanderaa C. QFeatures: Quantitative features for mass spectrometry data. [Internet]. 2024. Available from: https://github.com/RforMassSpectrometry/QFeatures.

13. Morgan M, Obenchain V, Hester J, Pagès H. SummarizedExperiment [Internet]. 2024. Available from: https://bioconductor.org/packages/SummarizedExperiment

14. Rainer J, Vicini A, Salzer L, Stanstrup J, Badia JM, Neumann S, et al. A Modular and Expandable Ecosystem for Metabolomics Data Annotation in R. Metabolites. 2022 Feb 11;12(2):173.

15. Kassambara A. rstatix: Pipe-Friendly Framework for Basic Statistical Tests [Internet]. 2023 [cited 2024 Apr 22]. Available from: https://cran.r-project.org/web/packages/rstatix/index.html

16. Ritchie ME, Phipson B, Wu D, Hu Y, Law CW, Shi W, et al. limma powers differential expression analyses for RNA-sequencing and microarray studies. Nucleic Acids Research. 2015 Apr 20;43(7):e47–e47.

17. Wickham H, Chang W, Henry L, Pedersen TL, Takahashi K, Wilke C, et al. ggplot2: Create Elegant Data Visualisations Using the Grammar of Graphics [Internet]. 2024 [cited 2024 Apr 22]. Available from: https://cran.r-project.org/web/packages/ggplot2/index.html

18. Plotly Technologies Inc. Collaborative data science [Internet]. Montréal, QC: Plotly Technologies Inc; 2015. Available from: https://plot.ly

19. Galili T, O’Callaghan A, Sidi J, Sievert C. heatmaply: an R package for creating interactive cluster heatmaps for online publishing. Wren J, editor. Bioinformatics. 2018 May 1;34(9):1600–2.

20. Colaert N, Helsens K, Martens L, Vandekerckhove J, Gevaert K. Improved visualization of protein consensus sequences by iceLogo. Nat Methods. 2009 Nov;6(11):786–7.

21. Wagih O. ggseqlogo: a versatile R package for drawing sequence logos. Hancock J, editor. Bioinformatics. 2017 Nov 15;33(22):3645–7.

22. Allaire JJ, Xie Y, Dervieux C, McPherson J, Luraschi J, Ushey K, et al. rmarkdown: Dynamic Documents for R [Internet]. 2024. Available from: https://github.com/rstudio/rmarkdown

23. Maturana FM. downloadthis: Implement Download Buttons in “rmarkdown” [Internet]. 2024. Available from: https://CRAN.R-project.org/package=downloadthis

24. Chang W, Cheng J, Allaire JJ, Sievert C, Schloerke B, Xie Y, et al. shiny: Web Application Framework for R [Internet]. 2024. Available from: https://CRAN.R-project.org/package=shiny

25. Rozanova S, Uszkoreit J, Schork K, Serschnitzki B, Eisenacher M, Tönges L, et al. Quality Control—A Stepchild in Quantitative Proteomics: A Case Study for the Human CSF Proteome. Biomolecules. 2023 Mar 7;13(3):491.

26. Cuervo AM. Cathepsin A regulates chaperone-mediated autophagy through cleavage of the lysosomal receptor. The EMBO Journal. 2003 Jan 2;22(1):47–59.

27. Yoon MC, Solania A, Jiang Z, Christy MP, Podvin S, Mosier C, et al. Selective Neutral pH Inhibitor of Cathepsin B Designed Based on Cleavage Preferences at Cytosolic and Lysosomal pH Conditions. ACS Chem Biol. 2021 Sep 17;16(9):1628–43.

28. Ivry SL, Sharib JM, Dominguez DA, Roy N, Hatcher SE, Yip-Schneider MT, et al. Global Protease Activity Profiling Provides Differential Diagnosis of Pancreatic Cysts. Clinical Cancer Research. 2017 Aug 15;23(16):4865–74.

